# Hydrogels improve plant growth in Mars analog conditions

**DOI:** 10.1101/2021.04.02.438167

**Authors:** Frédéric Peyrusson

## Abstract

The development of sustainable human presence in a Martian settlement will require *in situ* resource utilization (ISRU), the collect and utilization of Mars-based resources, including notably water and a substrate for food production. Plants will be fundamental components of future human missions to Mars, and the question as to whether Mars soils can support plant growth is still open. Moreover, plant cultivation will probably suffer from the lack of *in situ* liquid water, which might constitute one of the biggest challenges for ISRU-based food production on Mars. Enhancing the crop yield with less water input and improving water utilization by plants are thus chief concern for sustainable ISRU food production. Hydrogels are polymers able to absorb large quantity of water and have been shown to increase water retention in the soil, thus increasing plant establishment and growth. This work reports on the short-term assessment of plant growth in Mars soil analogs supplemented with hydrogels, in the constrained environment of a simulated life-on-Mars mission. Soil analogs consisted of sand and clay-rich material, with low amount organic matter and alkaline pH. Soils were supplemented with 10% (w/w) potting medium and were sampled in Utah desert, in the vicinity of the Mars Desert Research Station, a Mars analog facility surrounded by soils sharing similarities in mineralogical and chemical composition to Martian soils. Heights and dry biomass of *Mentha spicata* and seed germination of *Raphanus sativus* were monitored under various irrigation frequencies. Results indicate that the soil analogs, together with the limited irrigation regime, were less capable of supporting plant growth as a comparison to potting medium. The effects of hydrogel supplementation were significative under limited irrigation and led to growth increased by 3% and 6% in clay- and sand-containing soils, respectively. Similarly, hydrogel supplementation resulted in plant masses increased by 110% in clay-containing soils and 78% in sand-containing soils. Additionally, while seeds of *Raphanus sativus* failed to germinate, hydrogel supplementation allows for the germination of 27% of seeds, indicating that hydrogels might help loosening dense media with low water retention. Collectively, the results suggest that supplementation with hydrogel and traditional plant growth substrate could help plant cope with limited irrigation and poor alkaline Mars soil analogs, and are discussed in the context of strategies to develop ISRU for off-world colonization.

## INTRODUCTION

Sustained human settlement on Mars will require the use of plant-based bioregenerative advanced life support systems (ALS), with the potential to provide sustainable food production, air and water recycling, and to allow for the minimization of resupply missions (Richards et al. 2006; Ming 1989). Critically, such systems may likely depend on *in situ* resource utilization (ISRU), i.e., the use of existing materials at the settlement site, including notably water and a substrate for food production (Wheeler 2010). Together, ISRU and addition of ALS systems to exploration missions might save cargo volumes in spacecrafts, minimize safety issues, extend the length of planetary explorations and support mission success (Richards et al. 2006).

Due to the necessity of developing ISRU food production, the question as to whether plants can grow on Martian soils is of chief importance (Wamelink et al. 2014; Fox-Powell et al. 2016). In view of fleets of orbital and landed spacecraft, our understanding of Martian soils has improved considerably over the last few decades (Billi et al. 2019). Although Martian soils contain a variety of necessary micro- and macronutrients in accessible forms for plants (Ehrenfreund et al. 2011; Cannon et al. 2019), substantial soils properties argue against efficient plant growth, such as high concentrations of calcium perchlorate (Hecht et al. 2009), soils with low water retention capacity (Wamelink et al. 2014; Fox-Powell et al. 2016; Hecht et al. 2009), or soils moderately to highly alkaline (Hecht et al. 2009; Fairén 2008), a barrier for many plant species (Wamelink et al. 2005). In addition, the availability of *in situ* liquid water will be very limited on Mars (Bullock et al. 2004; Möhlmann 2004), water stress being one of the major factors limiting crop growth and plant biomass production (Shormin 2009). The study of Terrestrial analogs of Martian regolith on plant growth in the context of limited water resources are thus critically needed.

The Mars Desert Research Station (MDRS), operated by the Mars Society, is a Mars analog facility in Utah that allows Earth-based research to address the issues raised by space exploration, in the constrained environment of simulated life-on-Mars missions. MDRS is surrounded by a landscape that is an actual geologic Mars analog, with a mineralogy comparable to Mars, consisting of deposits of sands, clay minerals, iron oxides and traces of carbonates (Kotler et al. 2011; Direito et al. 2011).

Hydrogels are polymers able to absorb large quantities of water and fixing on seedling roots. By improving water availability, they have been shown to reduce water stress and improve plant growth and survival (Montesano et al. 2015). However, some studies report negative effects on different soil types (Del Campo et al. 2011), and no study has been conducted so far with Mars soil analogs. This study reports on the effect of hydrogels on plant growth and seed germination in two Mars soil analogs. All experiments were conducted during the UCL to Mars 2018 campaign within the constraints of *in situ* operations in the MDRS station (Saint-Guillain 2019; Wuyckens et al. 2019). Results demonstrate that hydrogel supplementation improves the growth of plants, despite poor and alkaline soils, when monitoring height and dry biomass of *Mentha spicata*. These effects were more pronounced in Mars soil analogs under limited irrigation. Hydrogel supplementation also improved the germination of *Raphanus sativus* seeds in dense sand-containing soils. The present work suggests that hydrogels could represent an interesting strategy for the use of *in situ* Mars soils.

## MATERIAL AND METHODS

### Soils sampling

Soil samples were collected in the vicinity of MDRS station, at an average altitude of 1391m, and consisted in a white sand layer and a brown-reddish clay-rich material (hereafter referred as “sand” and “clay”, respectively) (**Table 1**). These soils were previously analyzed for their composition during the EuroGeoMars 2009 campaign (Ehrenfreund et al. 2011), including elemental composition of nitrate, potassium, phosphorous, organic matter and carbonates. Soils were selected for their similarities with Mars soils, based on their (i) mineralogy (i.e., sand and phylosilicates [clay minerals]) and (ii) composition (i.e., low amount of organic matter, iron oxides, and traces of carbonates) (Boynton et al. 2009; Poulet et al. 2005; Chevrier and Mathé 2007). Additionally, soils were analyzed *in situ* for their pH in water, indicating pH of 9.2 and 9.06 for sand and clay soils, respectively (**Table 2**).

**Table 1.** Characteristics of soils used in this study. * Soils sampled at the Mars Desert Research Station location, adapted from (Ehrenfreund et al. 2011). N/A, not applicable or not available.

| Soil nature | Location | Organic matter (%) | N-element (ppm) | P (ppm) | K (ppm) | Carbonates (%) |
| --- | --- | --- | --- | --- | --- | --- |
| Sand * | N38.40746° W110.79280° | 1 | > 75 | 12 | 175 | < 1 |
| Clay * | N38.42638° W110.78342° | 2 | > 75 | 100 | 200 | 2 |
| Potting medium | N/A | 48 | < 10 | 22 | 150 | N/A |

**Table 2.**
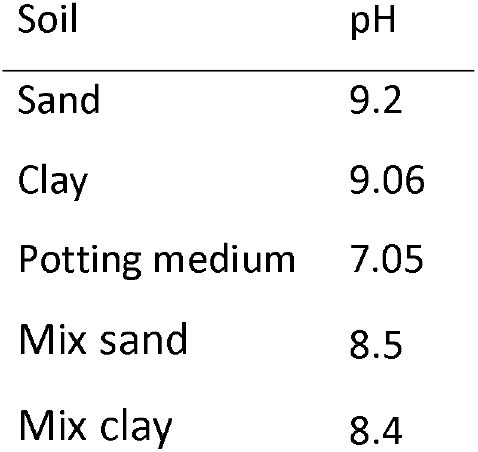
Determined pH of soils and mixed soils used in this study.

| Soil | pH |
| --- | --- |
| Sand | 9.2 |
| Clay | 9.06 |
| Potting medium | 7.05 |
| Mix sand | 8.5 |
| Mix clay | 8.4 |

### Water holding and release properties of hydrogels

Hydrogels are polymers that are able to increase water retention, and enhance plant growth (Montesano et al. 2015; Wang and Boogher 1987; El-Asmar et al. 2017). The superabsorbent polymer used in the present study was the commercial hydrogel STOCKOSORB^®^ 660 Medium (Evonik Industries; hereafter referred as “hydrogel”), a crosslinked potassium polyacrylic acid designed to remain active in the soil for 1 to 3 years. Soils were supplemented with 0.1% (w/w) hydrogel (according to the manufacturer’s instructions) prior to plant transplantation. Hydrogels were first analyzed for their ability to regenerate after drying, to mimic absorption by plant roots. A sample of 2 g of dry hydrogel was saturated with water, incubated 10 min prior to weighing. This sample was dried on absorbing paper at room temperature for 24 h, and subjected to three additional hydration/dehydration cycles.

### Experimental design for plant growth

All experiments were conducted in the greenhouse facility of MDRS station during the UCL to Mars 2018 campaign (Université catholique de Louvain), from Mars 12, 2018 (Sol 0) to Mars 25, 2018 (Sol 13), in a controlled atmosphere with an average temperature of 24 °C during the experimental period. During this period, mean length of visible light was 14 hours.

### Effect of hydrogels on plant growth

For a short-term assessment of plant growth, *Mentha spicata* plants were selected for their rapid growth and their robustness under nutrient-poor conditions. Experiments aimed at addressing the effect of hydrogel amendment in Mars soil analogs, in normal or reduced irrigation conditions. To that end, experiments were divided into three groups of soils: potting medium (“pot. medium“), sand supplemented with 10% (w/w) potting medium (“mix sand”), and clay supplemented with 10% (w/w) potting medium (“mix clay”). Each soil group was tested under normal or reduced irrigation regimes, performed in three independent experimental units. For each condition, seedlings of *Mentha spicata* were grown with or without 0.1 % (w/w) hydrogel supplementation. This experimental set-up resulted in three soils x two irrigation conditions x two supplementation conditions (with or without hydrogel supplementation) x three replicates.

Seedlings of *Mentha spicata* were transplanted at Sol 0 in single pots of 9 cm length and 10 cm depth with drainage outlet. Plants were manually overhead irrigated with water, daily for normal irrigation conditions, and each 4 days for reduced irrigation conditions, below requirements for seedlings of *Mentha* species (Shormin 2009; Clark R.J. 1980; McConkey et al. 2000). Shoot heights were recorded 13 days after transplantation (Sol 13). Initial seedlings were 23 ± 3 cm height. To normalize these variations and allow statistical comparisons, the growth of each plant was represented as a percentage of growth in comparison to its initial shoot height (Sol 0) (see **Figure 2A**).

Dry shoots masses were recorded for all plants 13 days after transplantation. Plants were dried in a forced air oven at 60 °C until constant mass (Valmorbida and Boaro 2007).

### Effect of hydrogels on seed germination

To monitor the first growth stages, seeds of *Raphanus sativus* were selected for their fast germination. Experiments were designed to address the effect of hydrogel-supplemented Mars soil analogs on seed germination. To that end, experiments were divided into five groups of soils: potting medium (“pot. medium”), sand supplemented with 10% (w/w) potting medium (“mix sand”), clay supplemented with 10% (w/w) potting medium (“mix clay”), sand and clay. This experimental set-up resulted in five soils x two supplementation conditions (with or without hydrogel supplementation) x three replicates. For each condition, five seeds were positioned per pot at the same burial depth. The percentage of seed germination were recorded at Sol 13.

### Statistical Analysis

Statistical analyses were performed with GraphPad Prism version 8.3.1, GraphPad InStat v3.10 (GraphPad Software) and SPSS v25.0 (IBM Statistics). For comparison between hydrogel and control conditions, statistical differences were determined using unpaired Student’s t-tests, with a threshold of statistical significance set to 0.05. *P* values strictly inferior to 0.05, 0.01 and 0.001 were used to show statistically significant differences and are represented with *, ** or *** respectively. For multiple comparison between soils, statistical differences were determined using one-way ANOVA with Tukey’s post hoc tests, with a threshold of statistical significance set to 0.05. Different letters indicate statistically significant differences.

## RESULTS

### Water holding and release properties of hydrogels

Hydrogels are polymers that can absorb large quantity of water during expansion (**Figure 1A**). When submitted to consecutives cycles of hydration and dehydration in order to mimic absorption by plant roots and watering, respectively, data indicated that they efficiently regenerate for sustained periods of time (**Figure 1B**). Additionally, roots of *Mentha spicata* transplanted in potting medium amended with 0.1% (w/w) hydrogel were able to fix on hydrogels (**Figure 1C**). These data indicate that hydrogels present interesting water holding and release properties that could constitute a method for improvement of plant growth.

**FIGURE 1.**
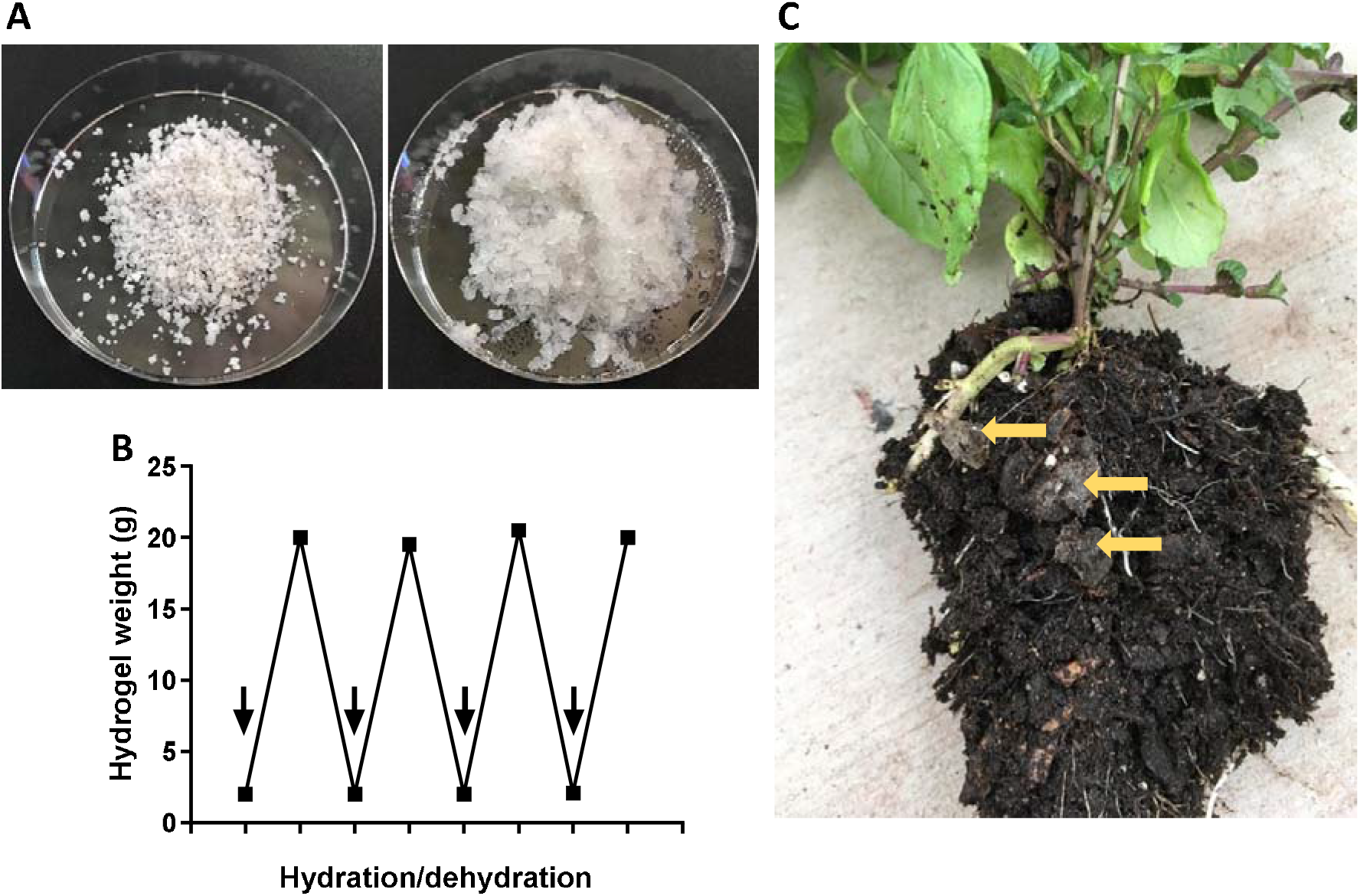
Water holding and release properties of hydrogels. (**A**) Sample of 1g of dry hydrogel (left). The same sample was saturated with water and incubated 10 min (right). (**B**) Hydrogels weight during hydration/dehydration cycles. A sample of 2 g of dry hydrogel was saturated with water (arrows), incubated 10 min prior to weighing, and dried on absorbing paper at room temperature for 24 h. (C) Hydrogels (arrows) fixed on *Mentha spicata* roots.

### Effect of hydrogels on plant growth

All experiments were conducted in the greenhouse facility of MDRS station from Mars 12, 2018 (Sol 0) to Mars 25, 2018 (Sol 13). For a short-term assessment of plant growth, *Mentha spicata* plants were selected for their rapid growth and their robustness under nutrient-poor conditions, and were allowed to grow for a total of 13 days post-transplantation. The mixing of soils with potting medium (groups mix sand and mix clay) decreased the initial pH from 9.2 to 8.5 and from 9.06 to 8.4, respectively (**Table 2**), and were in the range of pH of Martian soils (Fairén 2008), although bellow *Mentha* species requirements i.e., slightly acidic to neutral pH (Shormin 2009; Mohammadi and Asadi-Gharneh 2018; Valmorbida and Boaro 2007).

When grown in potting media, *Mentha* plants grew to 3% on average in 13 days, and the supplementation with hydrogels allowed for an 8% growth (**Figure 2A**), confirming the increased plant growth upon hydrogel amendment (Montesano et al. 2015). When compared to the control plants grown in potting medium without hydrogel supplementation, data indicated a significant reduction of plant growth in soil containing clay (101% their initial values; see uppercase letters in **Figure 2A**). By contrast, the growths recorded in clay- or sand- containing soils with hydrogel supplementation were statistically indistinguishable from those observed in potting medium (see lowercase letters in **Figure 2A**).

**FIGURE 2.**
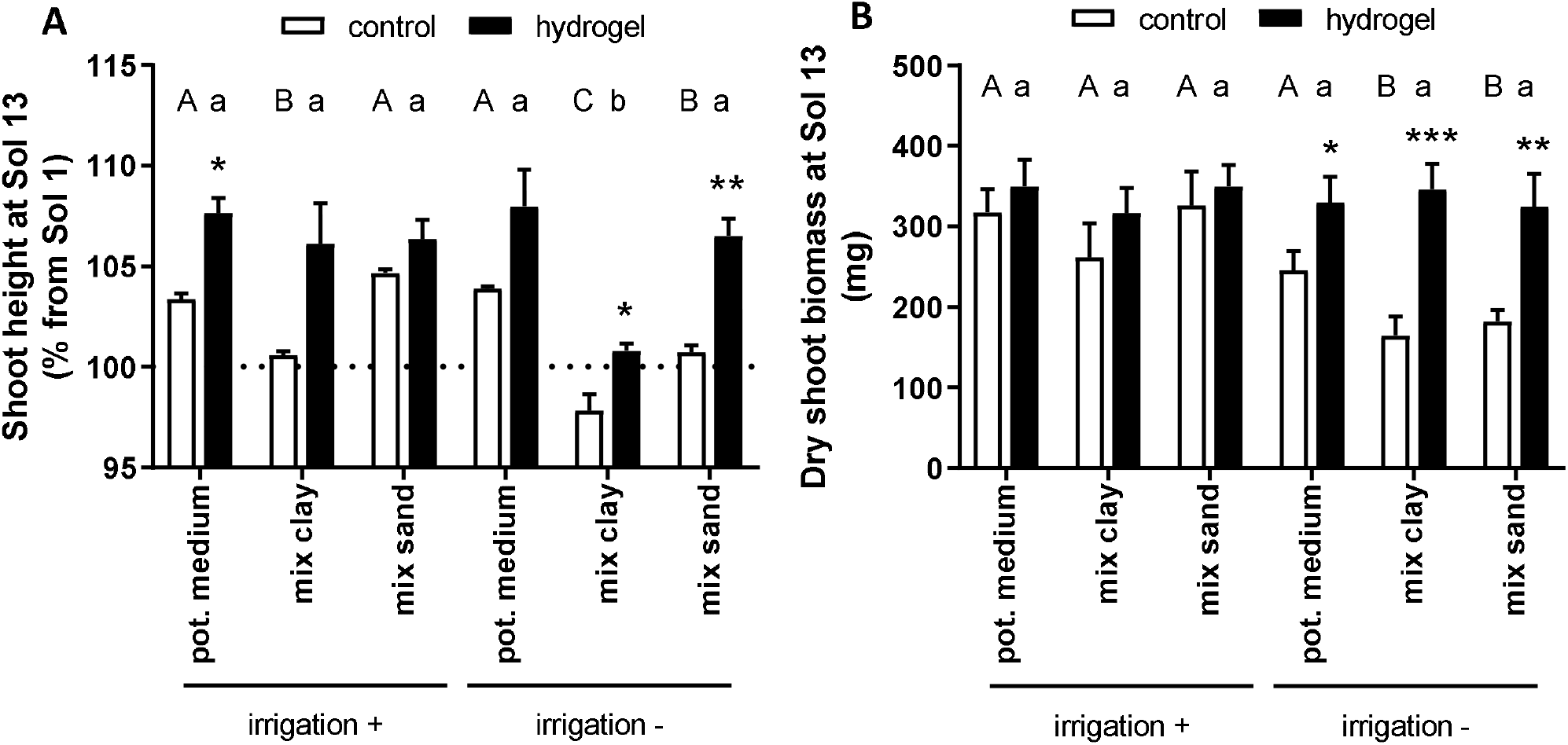
Shoots heights (**A**) and dry shoots biomasses (**B**) of *Mentha spicata* 13 days after transplantation (Sol 13), with (hydrogel) or without (control) 0.1 % (w/w) hydrogel supplementation, under varying irrigation conditions. Pots were filled with soils from MDRS location (see Table 1) supplemented with 10% (w/w) potting medium or with potting medium alone (mix clay, mix sand and pot. medium, respectively). Plants were either daily watered (irrigation +) or each four days (irrigation –). All data are means of three replications. Statistical analysis for comparison between control and hydrogel conditions: unpaired Student’s t-test. \**P* < 0.05, \*\**P* < 0.01, \*\*\**P* < 0.001. For comparison between soils: one-way ANOVA with Tukey’s post hoc test. Different uppercase (control) or lowercase (hydrogel) letters indicate values significantly different from each other.

Mint is a crop with a high water requirement during its active growth period (Shormin 2009). Due to the limited availability of liquid water for ISRU on Mars, the effects of hydrogel supplementation were then investigated under low irrigation conditions. To that end, plants were irrigated each four days, a low irrigation regime in comparison with requirements for *Mentha* seedlings (McConkey et al. 2000). When compared with potting medium without hydrogel supplementation, the limited irrigation led to reduced growth during the course of experiments, with plant heights of 97% and 101% their initial values, respectively (see uppercase letters in **Figure 2A**). These observations are consistent with reports showing that water stress significantly decreases plant height (Shormin 2009). Notably, a wilting of main branches of *Mentha* plants was observed in clay-containing soils, which appeared to be cohesive and unfavorable for plant growth, confirming that the deleterious effect of clay-containing soil was more pronounced under low irrigation frequencies.

Conversely, the growths recorded with hydrogel supplementation were not significantly affected among the different soils, except for clay-containing soil, albeit clay was able to support growth only in the presence of hydrogel. Under reduced irrigation frequencies, hydrogel amendment allowed for significantly improved growth in potting medium, clay- and sand-containing soils, leading to growth increased by 4%, 3% and 6% respectively, in comparison with non-amended conditions. These observations are in line with studies on *Argania spinosa* in arid region (C Defaa 2015), and on cucumber and basil plants in sandy soils (Montesano et al. 2015).

Similar trends were observed when monitoring biomasses, and the effect resulting from hydrogel amendment were significative under reduced irrigation (**Figure 2B**). In comparison with non-amended conditions, hydrogel supplementation resulted in plant masses increased by 34% in potting medium, 110% in clay-containing soils and 78% in sand-containing soils. Consistently, the plant masses recorded with hydrogel supplementation were not significantly different among the different soils or irrigation regimes.

Collectively, this suggests that supplementation with hydrogel and traditional plant growth substrate allowed for increased plant growth in poor alkaline Mars soil analogs and limited irrigation.

### Effect of hydrogels on seed germination

The effects of hydrogel supplementation in Mars soil analogs on seed germination was then assessed. To that end, seeds of *Raphanus sativus* were used due to their fast germination and were tested for their ability to germinate in five groups of soils: potting medium, sand or clay supplemented or not with potting medium.

All the seeds germinated in pots filled with potting medium regardless of the presence or absence of hydrogel amendment, indicating the absence of toxicity of hydrogels on seed germination in these conditions (Figure 3). A reduced germination was observed in soils containing clay, and sand to a lesser extent, allowing for the germination of 40 % and 66 % of seeds respectively, without significative effect of hydrogel supplementation. In control conditions, seeds of *Raphanus sativus* failed to show any germination after 13 days in pots only filled with clay or sand, which both create dense media largely unfavorable for germination. Similarly, clay did not support germination of seeds of *Raphanus sativus* under hydrogel supplementation.

**FIGURE 3.**
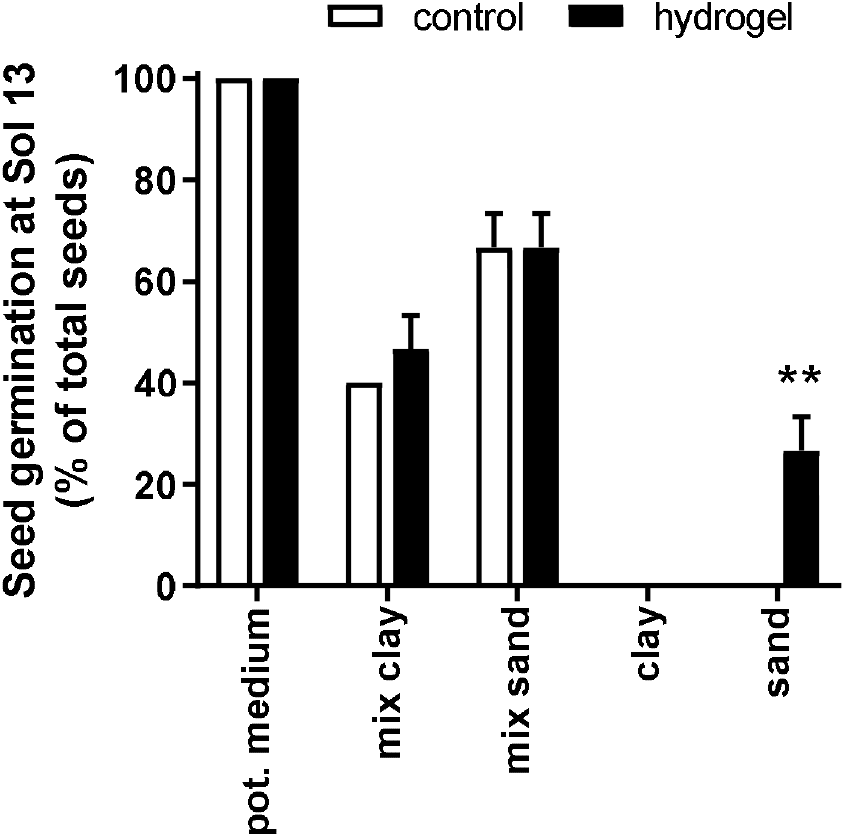
Percentage of seed germination of *Raphanus sativus* at Sol 13. For each condition, five seeds were positioned per pot, filled with potting medium, or soils from MDRS location (see Table 1) supplemented with 10% (w/w) potting medium, or soils alone, respectively, with (hydrogel) or without (control) 0.1 % (w/w) hydrogel supplementation. Data are means of three replications. Statistical analysis for comparison between control and hydrogel conditions: unpaired Student’s t-test. \*\**P* < 0.01.

Although sand is considered unfavorable to seed germination due to low water retention (Ren et al. 2002), hydrogel supplementation allows for the germination of 27% of seeds in average in sand. While preliminary, these results suggest that hydrogels could help loosening dense soils and facilitate seed germination in soils as the cohesive sand-like soils encountered in Mars (Arvidson et al. 2004a).

## DISCUSSION

Human missions to Mars will raise considerable number of challenges, among which the questions related to food self-sufficiency. Additionally, plants will be key components of bioregenerative advanced life support systems, as a potential source of food, and air, waste and water recycling.

This work takes place in body of studies that address the capacity of regolith simulants to support plant growth, in the context of ISRU for off-world colonization (Guinan 2018; Eichler et al. 2021; Wamelink et al. 2019; Wamelink et al. 2014). As a precursor to human visitations and future off-world agricultural systems, research regarding plant response to simulated Martian environment is of chief importance.

Our current knowledge of the Martian surface highly suggest that several mineral and chemical properties encountered will likely constitute barriers for plant growth. Firstly, Mars is covered with vast expanses of dust and coarse, sand-sized or finer-grained particles (Bishop et al. 2002). Sampling by Phoenix, and Spirit and Opportunity rovers revealed cohesive soils (Arvidson et al. 2004a; Arvidson et al. 2004b), and additionally, traces of clay minerals have been confirmed by Curiosity (Milliken et al. 2010). Low amounts of organic matter and carbon stored in carbonates have been found (Leshin et al. 2013; Boynton et al. 2009). Secondly, the measurements so far indicate that soils are alkaline, with pH ranging from 8 to 9 (Fairén 2008), which may be problematic for many plant species, by decreasing nutrients availability for plants (Wamelink et al. 2005). Finally, the availability of water for optimal plant cultivation might constitute one of the biggest issues raised by ISRU-based food production systems on Mars. Although liquid water has been evidenced, its hypersalinity might be a major limiting factor for its *in situ* use (Fox-Powell et al. 2016).

Several studies addressed the growth of plants in Martian regolith simulants, but some display critical differences with the chemical or mineralogical composition of Martian regolith such as the presence of organic matter (Seiferlin et al. 2008), or pH below the alkaline pH recorded at the Phoenix lander site (Hecht et al. 2009; Fairén 2008).

The proposed study tends to integrate these different parameters, and to assess the effect of hydrogel on plant growth in Mars soil analogs, consisting in sand and clay-rich material with low amount organic matter and alkaline pH.

Although experiments were limited to growth indices of *Mentha spicata*, results indicate that these soils, together with the limited irrigation frequency, were less capable of supporting plant growth as a comparison to potting medium. This indeed resulted in culture conditions well beyond *Mentha* species requirements i.e., slightly acidic pH media (Shormin 2009; Mohammadi and Asadi-Gharneh 2018; Valmorbida and Boaro 2007) and frequent irrigation (Shormin 2009; Clark R.J. 1980; McConkey et al. 2000). Under water stress, N-uptake as well as vegetative growth and biomass production of plants significantly decrease, leading to more pronounced effect of hydrogel supplementation under limited irrigation. Indeed, hydrogel amendment allowed for increased plant heights and biomass in Mars soil analogs under low irrigation regime. Additionally, results indicate that soil analogs were unable to support seed germination of *Raphanus sativus*, while hydrogel supplementation allowed for seed germination in sand, and suggest that hydrogels could help loosening dense soils. Collectively, these data indicate that supplementation with hydrogel and traditional plant growth substrate could help plants cope with poor, alkaline Mars soil analogs and limited watering.

More globally, this work further confirms the need of soil supplementation outlined by several studies to support the growth of plants. This notably comprises supplementation with nitrogen (through direct NH_4_/NO_3_ supplementation or nitrogen fixing bacteria), an essential nutrient for plant growth, which is absent in JSC-Mars-1A simulant (Wamelink et al. 2019), although Curiosity Mars Science Laboratory detected nitrate in sedimentary and aeolian deposits within Gale crater (Stern et al. 2015). Other reports highlight the importance of soil acidification to improve the plant viability, and the necessity of detoxifying substrates from perchlorates (Eichler et al. 2021), through e.g., perchlorate-reducing bacteria (Coates John et al. 1999).

This work is a short-term assessment conducted within the constraints of *in situ* operations and will require further studies on other plants, Mars soil analogs and hydrogels. Together, these results should be considered as a proof of concept that indicate the potential interest of hydrogels to limit water input in ISRU-based food production systems.

## AUTHOR CONTRIBUTIONS

FP conceived and designed the experiment, collected the data, performed the analysis and wrote the paper.

## FUNDING

The author declares no competing interests. FP is recipient of a postdoctoral fellow from the *Université catholique de Louvain*.

## ACKNOWLEDGEMENTS

This work could not have been conducted without the involvement of the entire UCL to Mars 2018 crew: Bastien Baix, Martin Roumain, Michael Saint-Guillain, Ariane Sablon, Sophie Wuyckens, Mario Sundic and Maximilien Richald, who all contributed to a relevant and inspiring campaign. I also gratefully acknowledge the Mars Society, especially Robert Zubrin, Shannon Rupert and the Mission Support team, for creating the conditions that make rotations at the MDRS relevant to Mars analog sojourns.

## Notes

### Competing Interest Statement

The authors have declared no competing interest.

